# Cryo-EM structure of the vault from human brain reveals symmetry mismatch at its caps

**DOI:** 10.1101/2025.05.27.656403

**Authors:** Sofia Lövestam, Sjors H.W. Scheres

**Affiliations:** MRC Laboratory of Molecular Biology, Cambridge, UK

## Abstract

The vault protein is expressed in most eukaryotic cells, where it is assembled on polyribosomes into large hollow barrel-shaped complexes. Despite its widespread and abundant presence in cells, the biological function of the vault remains unclear. In this study, we describe the cryo-EM structure of vault particles that were imaged as a contamination of a preparation to extract tau filaments from brain tissue of an individual with progressive supranuclear palsy (PSP). We identify a mechanism of symmetry mismatch at the caps of the vault, from 39-fold to 13-fold symmetry, where two out of three monomers are sequentially excluded from the cap, resulting in a narrow, greasy pore at the tip of the vault. Our structure offers valuable insights for engineering carboxy-terminal modifications of the major vault protein (MVP) for potential therapeutic applications.

## Introduction

The vault, first identified in 1986 by electron microscopy (EM) as a contaminant in clathrin-coated vesicle preparations from rat liver, is an abundant, hollow, barrel-like macromolecular complex found in the cytoplasm of nearly all eukaryotic cells ^1,2^. Vault particles are among the largest known macromolecular complexes in the cell: they measure approximately 41 × 41 × 72 nm and have a molecular weight of ∼13 MDa. The structure of the shell of the vault was first solved by X-ray crystallography ^3^ and subsequently by cryo-EM single-particle analysis ^4,5^. It consists of two symmetrically packed half-vaults, each formed by 39 copies of the major vault protein (MVP), a 100 kDa protein that consists of 12 domains: nine N-terminal structural repeats (R1 to R9), a shoulder or SPFH domain, a cap helix, and a carboxy-terminal cap-ring domain. In both X-ray and cryo-EM structures ^3,4^, the application of 39-fold symmetry reduced the information available about the structure at the top and bottom caps of the vault, where there is insufficient space to accommodate 39 monomers.

The function of the vault remains unknown. The protein shell has been reported to encapsulate vPARP (193 kDa) ^6^, telomerase associated protein-1 TEP1 (240 kDa) ^7^, and a small untranslated RNA (vault RNA) of 141 bases^1^. The vault has been found to be expressed more highly in cancerous cells ^8,9^ and has been suggested to play a role as toxin remover. Elevated levels of MVP have implicated the vault in the development of multidrug resistance in (MDR) in various cancers ^10,11^.

The large size and non-immunogenic properties of the vault makes it an attractive vehicle for bioengineering approaches to delivering a diverse range of molecules into cells. The amino-terminus of the MVP is situated at the waist of the vault, where it embeds into the lumen (i.e. the interior of the vault); the carboxy-terminus extends outwards at the cap ^4^. Both termini have been engineered to facilitate targeted delivery and encapsulation of therapeutic compounds. Incorporating additional amino acids at the carboxy-terminus enables the display of targeting peptides on the vault surface, promoting specific interactions with cell surface receptors. For instance, recombinant vaults with engineered carboxy-terminal extensions have been shown to bind to epithelial cells directly, or via a monoclonal antibody targeting the epidermal growth factor receptor (EGFR) ^12^. Additionally, it has been shown that introducing an amphipathic α-helix at the amino-terminus of the MVP (i.e. in the lumen of the vault) creates a microenvironment that is suitable for encapsulating hydrophobic compounds and thereby enhances the delivery of lipophilic drugs into cells ^13^.

We regularly observe vaults particles in cryo-EM images of amyloid filaments that we extract from human brain tissue with neurodegenerative disease using sarkosyl solubilisation and ultracentrifugation. Here, we calculate a cryo-EM reconstruction using vault particle images in cryo-EM micrographs of brain extracts from an individual with progressive supranuclear palsy (PSP), which we had previously deposited in the EMPIAR database (EMPIAR-10765 ^14,15^). Using symmetry expansion and focused image classification techniques in RELION ^16^, we reveal how the 39-fold symmetry at the caps of the vault is broken by the sequential exclusion of two MVP monomers, leading to a narrow pore on each cap of the vault that comprises only 13 MVP monomers. This structure could be engineered for targeted drug delivery.

## Results

Cryo-EM micrographs of brain extracts from an individual with progressive supranuclear palsy (PSP)^15^showed multiple vault particles, most of which contained internal density and some of which were empty (**Figure 1A**). We predominantly observed intact vaults but occasionally also detected half-vaults, in accordance with previous reports ^4^.

**Figure 1.**
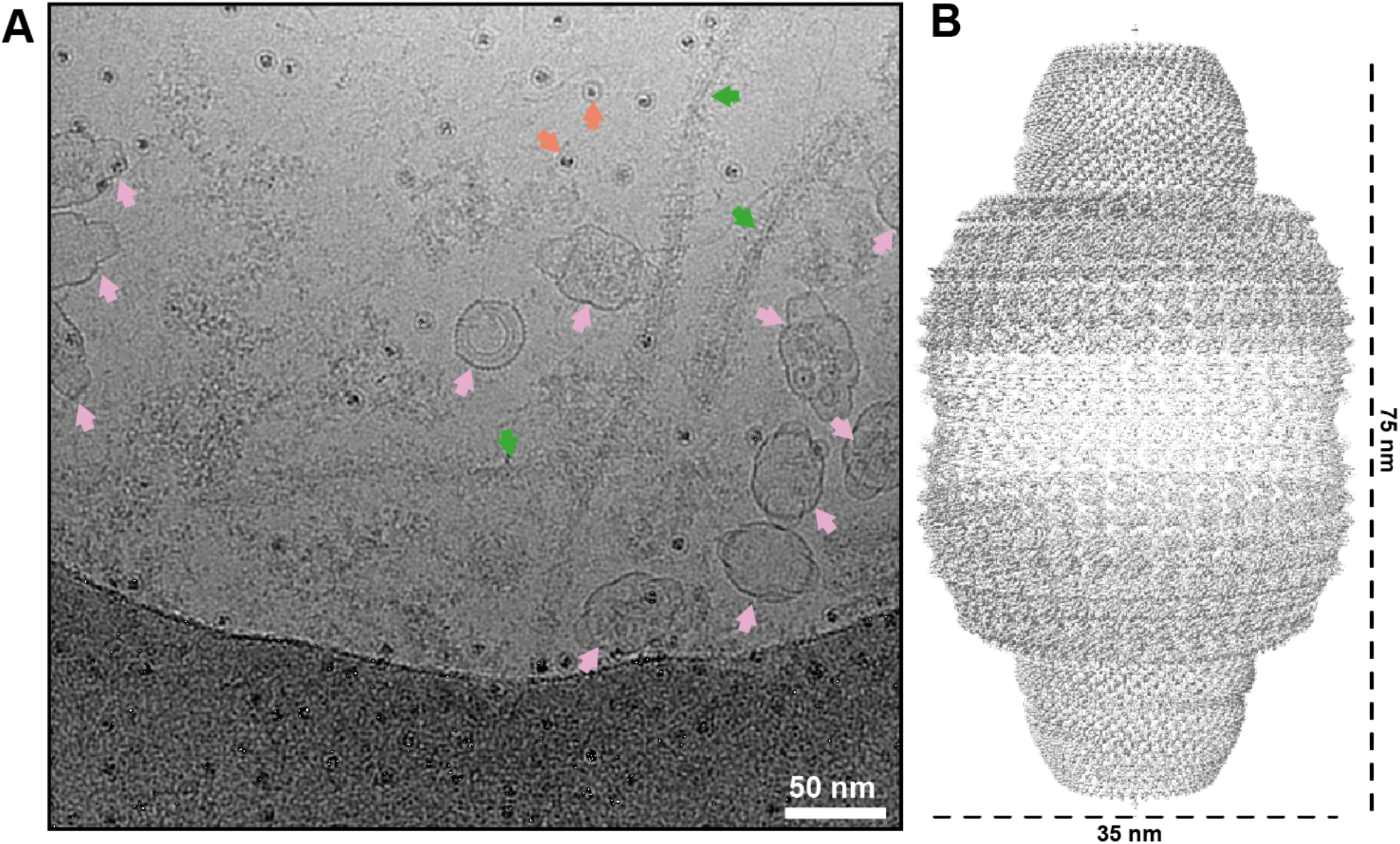
Cryo-EM of the vault particle. **A** Cryo-EM micrograph of vault particles. Pink arrows indicate vaults; green arrows are PSP filaments and orange arrows are ferritin particles. **B** Electron density map of the vault.

Using D39 symmetry, we determined the structure of the vault complex at an overall resolution of 3.1 Å (Methods**; Figure 1B**; **Extended Data Figure 1A**). Albeit at higher resolution, our D39 structure is nearly identical to previously reported structures of the vault in the primed state ^4^. We used the D39 map to build an atomic model for the majority of the MVP residues (11-803), which comprise the structural repeats (R1-R9), the shoulder domain, and the cap helix domain (**Figure 2**). Lower local resolution (4.0 Å) for the first three repeats (R1-R3) suggests the presence of unresolved structural heterogeneity at the waist of the vault. We did not observe density for the first eleven residues of MVP, nor for any molecules that might be encapsulated within the vault. As with previously reported structures of the vault, the reconstructed density at the caps was smeared, which precluded atomic modeling of the carboxy-terminal residues 801-893.

**Figure 2.**
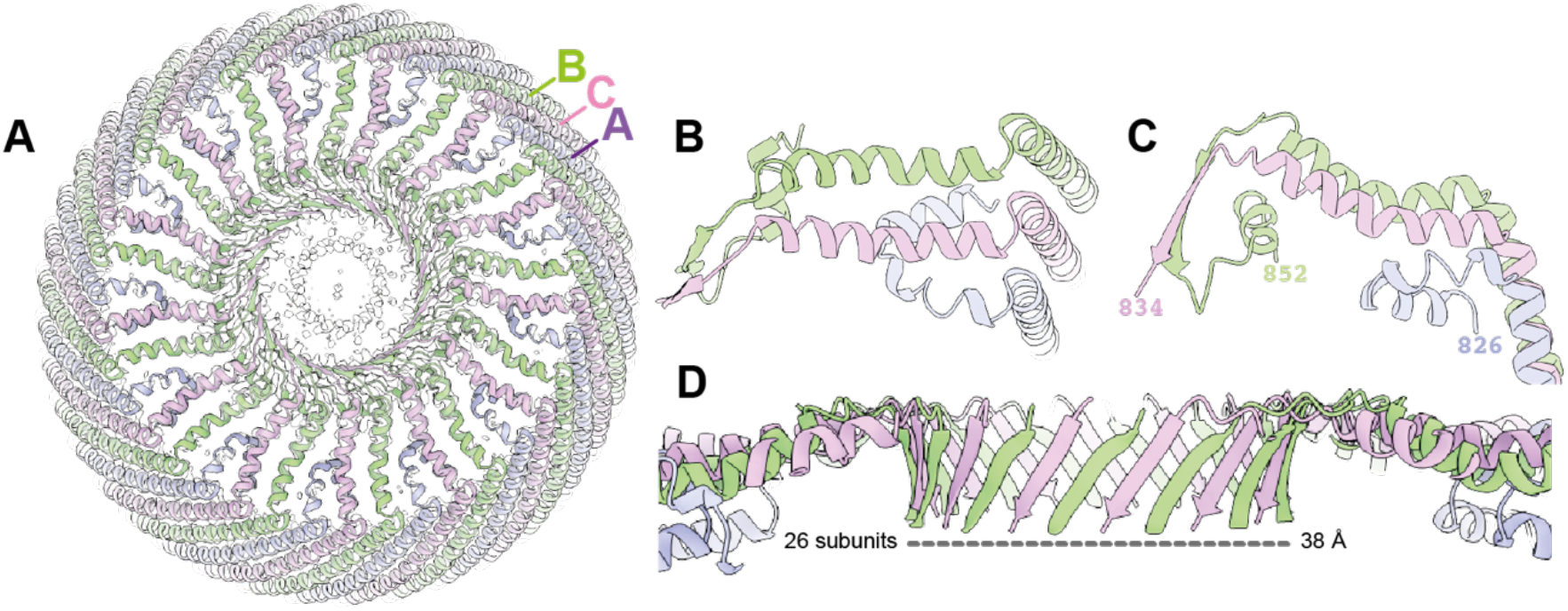
Cap of the vault. **A** Top view of the vault. Cryo-EM density is shown for the central region of the cap; a cartoon model is shown fitted into the density. **B** Top view of the pore showing the exclusion of monomer A (in purple) **C** As in B but lateral orientation. **D** Side view of the parallel β-barrel formed by 26 MVP subunits.

**Figure 3.**
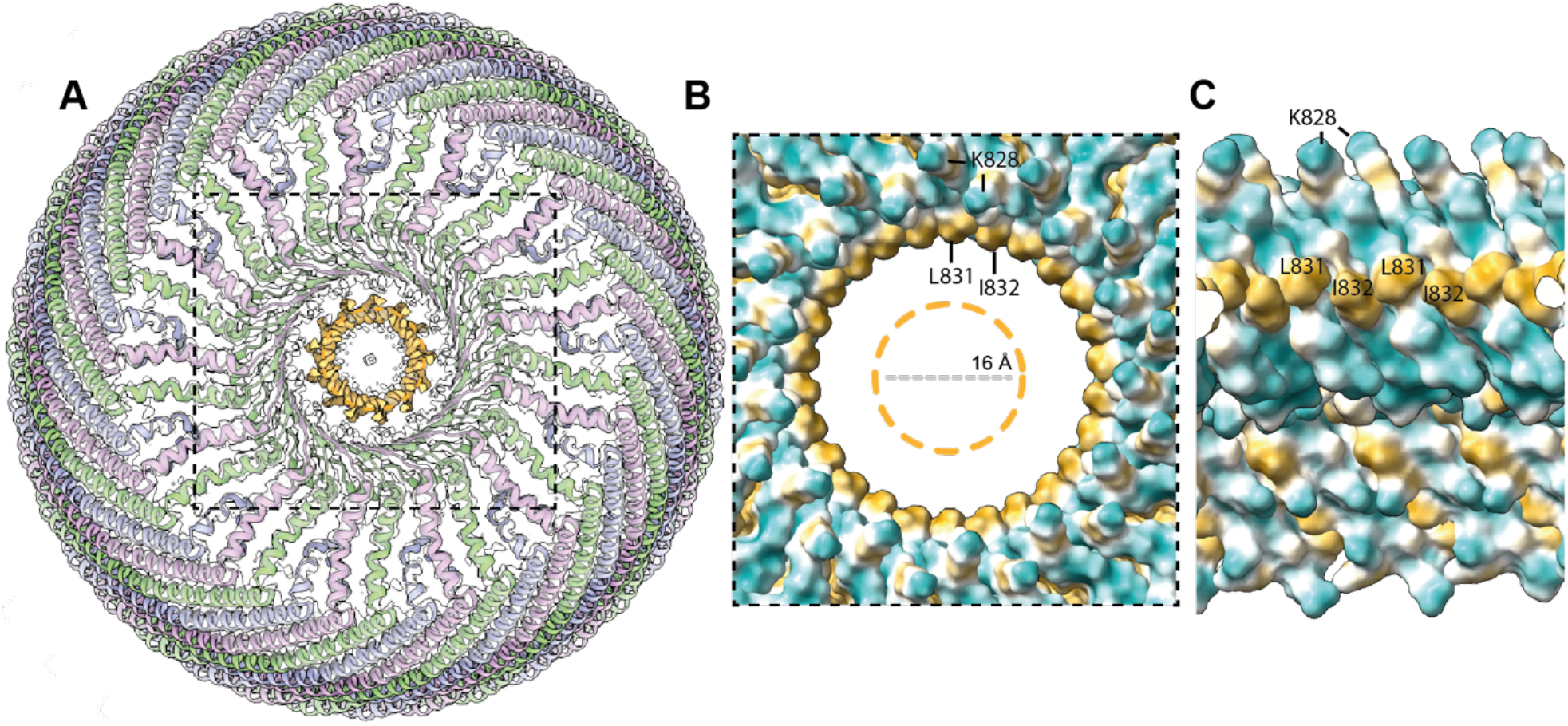
Hydrophobic features at the cap of the vault. **A** Top view of the vault model with unmodeled density highlighted in orange. **B** Hydrophobicity plot showing a hydrophobic ring made by residues L381 and I382 on the inside of the 26-membered parallel β-barrel. The central unmodeled density is indicated by an orange dashed circle. **C** Cross-sectional view of the pore hydrophobicity plot.

To improve the reconstructed density at the cap of the vault, we used D39 symmetry expansion, followed by two rounds of focused 3D classification in C13 symmetry on one of the caps (see Methods). This procedure resulted in a map with a resolution of 3.6 Å that shows three distinct arrangements of MVP monomers, where two out of every three MVP monomers are sequentially excluded from the cap structure.

In the monomer that is excluded first (monomer A; purple in Figure 2), the long α-helix that spans residues 803-823 in the other two monomers breaks at residue 813, and the chain folds towards the lumen, where residues 815-826 form another short α-helix. In the other two monomers (monomers B and C; green and pink in Figure 2), following the long α-helix that comprises residues 803-823, the chains transition into β-strands that comprise residues 829-834 and that form a 26-membered parallel β-barrel, with a diameter of 38 Å and with all chains pointing down towards the centre of the vault.

The monomer that is excluded second (monomer B; green in Figure 2), then forms a small loop and a short α-helix, comprising residues 835-853, that packs against the outside of the 26-membered parallel β-barrel, and below the long α-helices formed by residues 803-823 of monomers B and C.

Finally, at the centre of the cap, the cryo-EM density becomes too smeary to identify the position of residues beyond D834 for monomer C. However, weaker density in this region suggests that the carboxy-terminal residues of monomer C may fold back upwards, forming what is possibly a second parallel β-barrel-like pore with 13 β-strands and a diameter of 16 Å. It is possible that the weak cryo-EM density in this central pore is a consequence of variations in the specific residues that make up this pore. Based on the amino acid sequence of MVP, residues at both the inside and outside of this barrel are likely to be of a hydrophobic nature. This greasy pore may adopt multiple orientations with respect to the wider 26-membered parallel β-barrel, which also forms a hydrophobic ring made by residues L831 and I832 (**Figure 4**).

**Figure 4.**
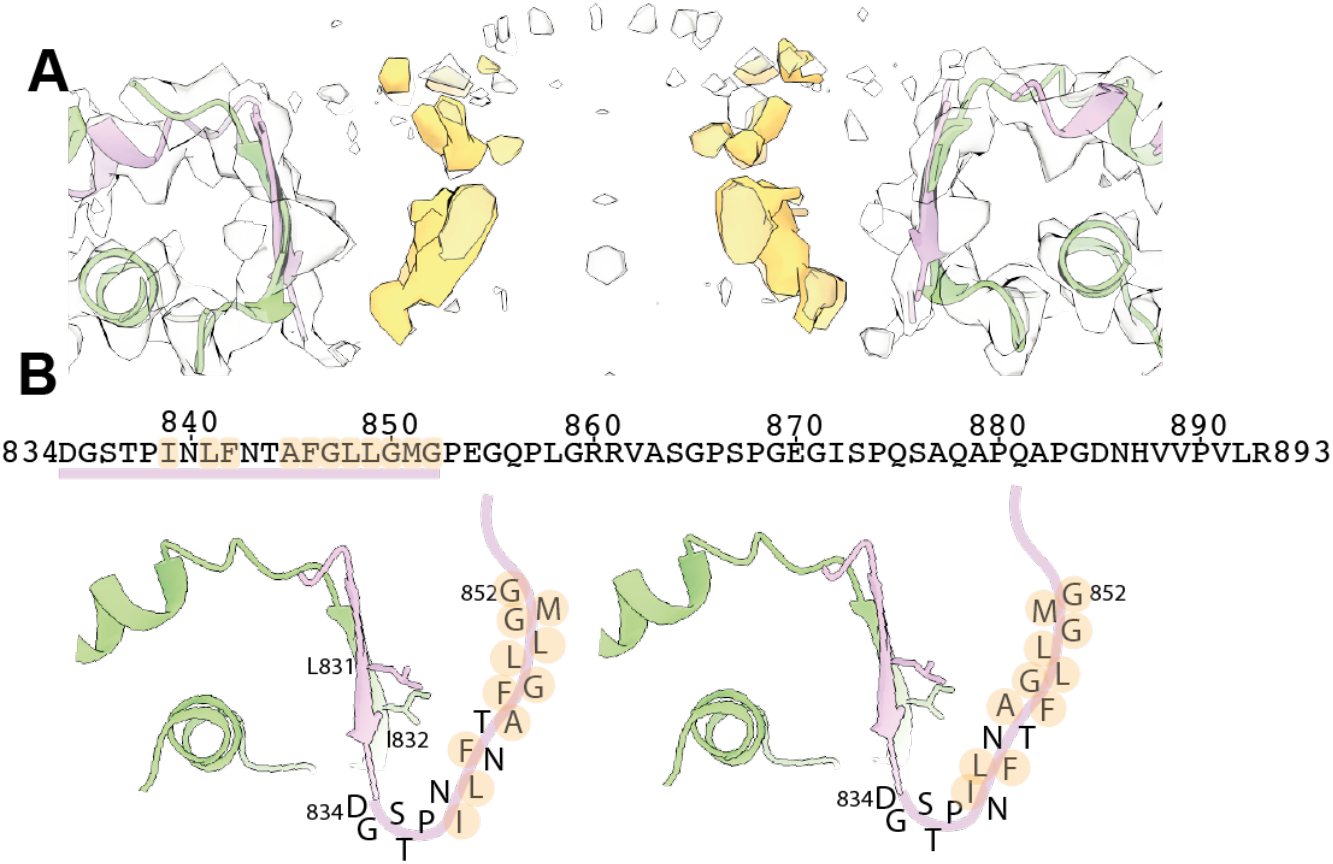
Structure and sequence of the 13-subunit pore. **A** Cross-sectional view of the central portion of the vault cap. The vault structure is shown as a cartoon, and the unmodelled density is shown in orange. **B** Primary sequence of the vault spanning residues 834-893. Putative residues which line the pore are shown with their one-letter code in two possible arrangements. Hydrophobic residues are highlighted in orange.

## Discussion

Although the vault is among the largest known protein complex, and MVP is of high abundance and conservation in eukaryotes, its function remains a mystery. Nevertheless, a better understanding of the structure of the vault is of importance for engineering these complexes for the delivery of therapeutics, as encapsulation of cargo within the vault’s protein shell could potentially bypass the immune system for the delivery of immunogenic therapeutics ^17^.

Previous studies were unable to resolve the cap of the vault and instead hypthesised that the monomers cross over and form a double-layered cap to accommodate the 39 MVP chains ^18^. Our cryo-EM reconstruction now reveals that the 39-fold symmetry is broken by the sequential exclusion of two out of every three monomers from the cap structure. Our density for the region closest to the central axis of the vault still does not support atomic modeling, probably due to remaining conformational heterogeneity that we were unable to resolve in our image processing. Weaker density in this region of the map suggests that an inner pore is made by a 13-stranded β-barrel. A comparable mechanism has been observed in the Erlin1/2 complex, which is composed of 13 subunits of Erlin1 and 13 subunits of Erlin2, forming a 26-mer “half-vault” barrel that sits on the membrane of the endoplasmic reticulum. At the cap of the Erlin1/2 complex, Erlin2 is selectively excluded, resulting in a 13-stranded β-barrel formed exclusively by Erlin1^19^. Given the hydrophobic nature of the residues in the corresponding part of the MVP sequence, the 13-stranded β-barrel at the vault cap is likely to have a greasy interior, which would favour the passing of hydrophobic molecules, or preclude the passing of polar or charged molecules. Modification of this part of the MVP sequence to residues of a different chemical nature could be used to make this pore selective for different types of molecules entering the vault. Thereby, while questions regarding the function of the vault remain to be deciphered, our structure will inform bioengineering approaches that seek to alter which molecules pass through its cap, or approaches that aim to place signals on the cap that target it towards specific cell types. Furthermore, the mechanism of symmetry mismatch identified in this study implies that any modification to the carboxy-terminal residues of MVP will remain inside the lumen of the vault for two out of every three monomers.

## Methods

### Cryo-EM image processing

Cryo-EM micrographs were downloaded from EMPIAR^14^ (accession code 10765^20^) and processed in RELION^21^. Movies were motion corrected using RELION’s implementation of MotionCor2, and Contrast Transfer Function (CTF) parameters were estimated using CTFFIND-4.1^22^. Manually picked vault particles from 50 micrographs were used to train a Topaz ^23^ model to pick the remaining 3561 micrographs. Auto-picked particles were extracted in a box size of 640 Å and a pixel size of 5.75 Å. Reference-free 2D classification was used to assess the quality of the picking and showed the presence of views from multiple directions. Selecting 2D class averages with vault particles, we re-extracted 17,202 particles with a box size of 735.8 Å and a pixel size of 1.30 Å. Subsequent 3D auto-refinement in D39 symmetry and starting from an angular sampling rate of 1.8 degrees was performed using a published vault cryo-EM map (EMDB 13478^20,24^), low-pass filtered to 60 Å, as an initial model. 3D classification was used to remove suboptimal particles, resulting in a data set of 7,367 particles that were used for a final 3D refinement with D39 symmetry and CTF refinement ^25^. Standard post-processing procedures in RELION were used to obtain a sharpened map with a resolution of 3.1 Å.

To improve the reconstructed density at the cap of the vault, we used D39 symmetry expansion of the particles from the final 3D auto-refinement, resulting in 574,626 expanded particles. We then performed focused 3D classification with partial signal subtraction ^26^ of the rest of the vault density on one of the caps, with ten classes, imposing C13 symmetry, and without further optimisation of the particle poses from the refinement in D39 symmetry. We selected 129,211 expanded particles that belonged to two of the classes that showed clear structures with 13-fold symmetry at the centre of the cap. The map of the second class was rotated -360°/39 with respect to the map of the first class. The remaining 445,415 particles were subjected to a second round of 3D focused classification into ten classes, which identified another 48,584 particles that showed clear structures with 13-fold symmetry at the centre of the cap. This map was rotated +360°/39 with respect to the map of the first class from the first round of focused classification. Two half-maps of the cap structure were then reconstructed using the relion_reconstruct program from the original half-sets of the combined 177,795 expanded particles, imposing C13 symmetry and using the original poses from the 3D refinement with D39 symmetry (while modifying the first Euler (rot) angles of the two rotated classes by +/-360°/39). Standard RELION post-processing yielded a final map of the cap structure to an estimated resolution of 3.6 Å.

### Model building

The model in the map with D39 symmetry was rebuilt from PDB entry 4hl8^4^. The model for the cap of the vault was built using ModelAngelo ^27^. Both models were refined in ISOLDE ^28^. The map and model with D39 symmetry and for the cap of the vault were deposited in the PDB and EMDB and are available under accession codes 9R86, 9R87 and 53805, 53806, respectively.

## Acknowledgements

We thank Max Wilkinson and David Li for their enthusiasm surrounding the mysteries of the vault protein; Toby Darling, Ivan Clayson and Jake Grimmett for assistance with high-performance computing; and Leonard Rome for critical reading of the manuscript. S.L. is grateful to Anne Bertolotti for encouragement to work on proteins that are not related to her project. This work was supported by the MRC, as part of the U.K. Research and Innovation (UKRI) (MC_UP_A025_1013 to S.H.W.S.).

## Author contributions

S.L and S.H.W.S. both analysed cryo-EM data and jointly wrote the manuscript.

---

**Extended Data Figure 1.**
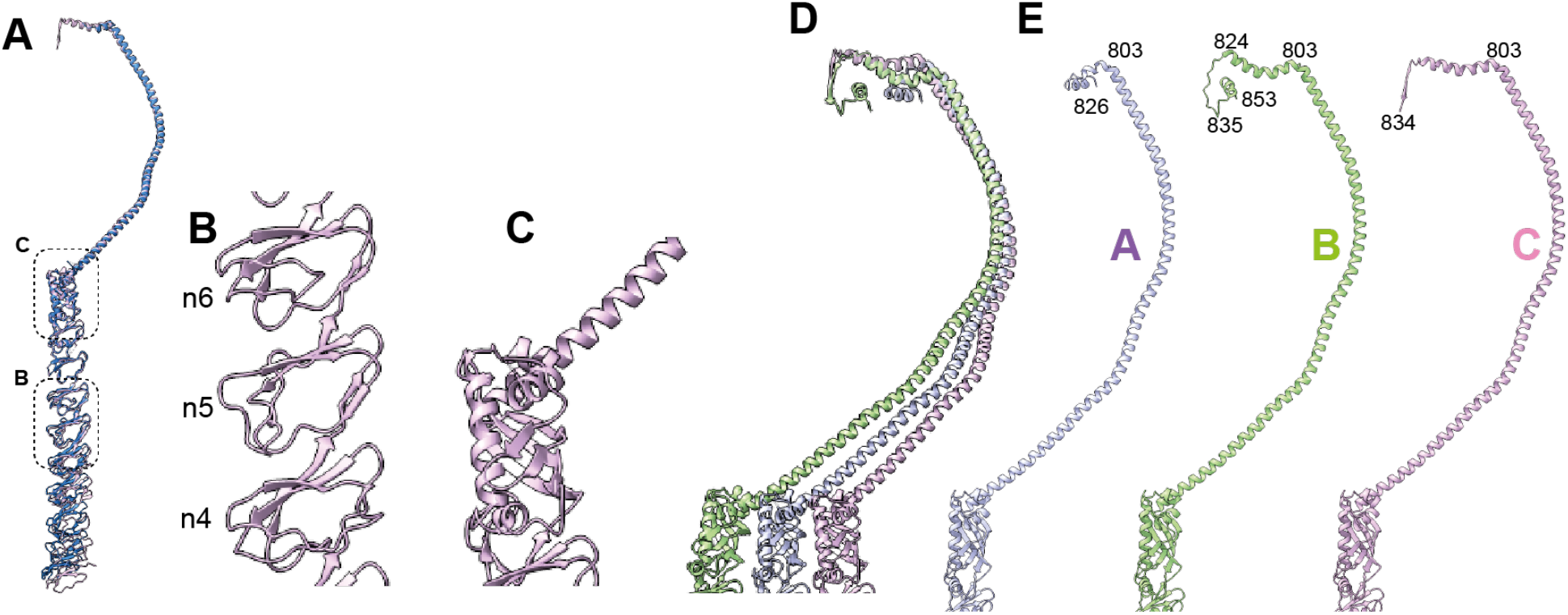
Structure of the Major Vault Protein subunit. **A** Major Vault Protein (MVP) from our cryo-EM structure (purple) overlaid with previously solved MVP structure (PDB:4hl8 in blue)^4^. **B** Close-up of n4, n5 and n6 repeat regions of the MVP **C** Close-up of the shoulder domain **D** Close-up of three different arrangements of the carboxy-terminus of the vault monomers where A is shown in purple, B in green and C in pink. **E** Separate views of the carboxy-terminus of monomers A-C.

**Extended Data Figure 2.**
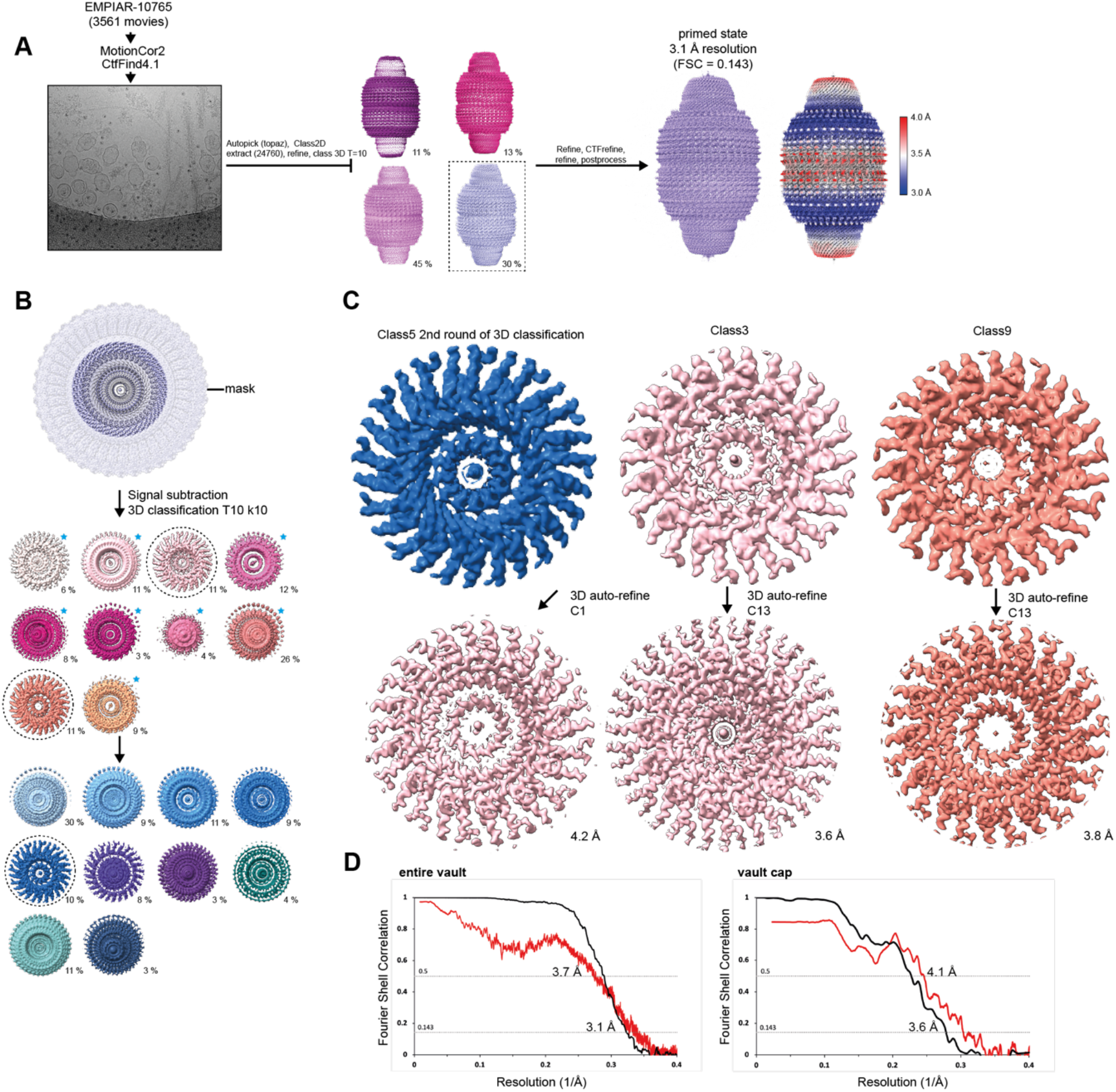
Cryo-EM data processing. **A** Cryo-EM micrographs were imported, corrected for motion and CTF parameters were estimated. Particles were autopicked using Topaz and particles were refined in RELION. 3D classification was used to select the best particles for CTF-refinement and for high-resolution 3D auto-refinement. Local resolution of the post-processed map is coloured from blue (3.0 Å) to red (4.0 Å). **B** Processing of the cap of the vault. A mask was made to include the cap of the vault and use partial signal subtraction for the remainder of the vault density. 3D classification using C13 symmetry was used to identify and select the best particles. **C** Selected particles were corrected for their first Euler angles by +/-360°/39 and used for 3D reconstruction in C13 symmetry, followed by standard RELION post-processing for resolution estimation and sharpening. **D** Fourier Shell Correlation curves are shown for two independently refined cryo-EM half maps (black) and for the final refined atomic model against the final cryo-EM map (red) for the D39-symmetric map of the entire vault and the C13 symmetric map of the cap.

**Table 1:** Model validation.

| **Data processing** | **Entire vault**<br>**PDB: 9R86**<br>**EMDB: 53805** | **Vault cap**<br>**PDB: 9R87**<br>**EMDB: 53806** |
| --- | --- | --- |
| Initial particle images (no.) | 24760 | 574626 |
| Final particle images (no.) | 7367 | 65387 |
| Symmetry imposed | D39 | C13 |
| Map resolution FSC 0.143 (Å) | 3.1 | 3.6 |
| **Refinement** |  |  |
| Initial model used (PDB code) | 4hl8 | ModelAngelo |
| Model resolution FSC 0.5 (Å) | 3.7 | 4.1 |
| Map sharpening *B* factor (Å<sup>2</sup>) | -76 | -98 |
| Model composition |  |  |
| Non-hydrogen atoms | 220,580 | 20,605 |
| Protein residues | 27,651 | 2704 |
| Ligands | na | na |
| *B* factors (Å<sup>2</sup>) |  |  |
| Protein | 95.61 | 96.82 |
| Ligand | na | na |
| R.m.s. deviations |  |  |
| Bond lengths (Å) | 0.011 | 0.010 |
| Bond angles (°) | 1.932 | 1.873 |
| Validation |  |  |
| MolProbity score | 0.74 | 0.84 |
| Clashscore | 0.13 | 0.07 |
| Poor rotamers (%) | 0.25 | 0.62 |
| Ramachandran plot |  |  |
| Favored (%) | 96.89 | 95.58 |
| Allowed (%) | 3.11 | 4.30 |
| Disallowed (%) | 0 | 0.11 |

